# Human Demographic Outcomes of a Restored Agro-Ecological Balance

**DOI:** 10.1101/637777

**Authors:** K.A.G. Wyckhuys, D.D. Burra, J. Pretty, P. Neuenschwander

## Abstract

As prominent features of the Anthropocene, biodiversity loss and invasive species are exacting serious negative economic, environmental and societal impacts. While the monetary aspects of species invasion are reasonably well assessed, their human and social livelihood outcomes often remain obscure. Here, we empirically demonstrate the (long-term) human demographic consequences of the 1970s invasion of a debilitating pest affecting cassava -a carbohydrate-rich food staple-across sub-Saharan Africa. Successive pest attack in 18 African nations inflicted an 18 ± 29% drop in crop yield, with cascading effects on birth rate (−6%), adult mortality (+4%) and decelerating population growth. The 1981 deliberate release of the parasitic wasp *Anagyrus lopezi* permanently restored food security and enabled parallel recovery of multiple demographic indices. This analysis draws attention to the societal repercussions of ecological disruptions in subsistence farming systems, providing lessons for efforts to meet rising human dietary needs while safeguarding agro-ecological functionality and resilience during times of global environmental change.

## Introduction

The UN Sustainable Development Goals (SDGs) pursue a global alleviation of malnutrition and poverty, improved human well-being and a stabilization of the Earth’s life-support systems (Griggs et al., 2013). Biodiversity lies at the core of multiple SDG targets, and -aside from its high intrinsic value-underpins the delivery of myriad ecosystem services (Wood et al., 2018). Yet, efforts to meet dietary requirements of a growing global population have negatively impacted upon biodiversity and the public goods supplied by natural ecosystems (Godfray et al., 2010; Fischer et al., 2017). Land conversion, ecosystem mis-management and the negative externalities of agricultural intensification continue to wage a toll on the world’s biodiversity (Maxwell et al., 2016; Isbell et al., 2017; Pretty et al., 2018). These human-mediated processes risk destabilizing both terrestrial and marine ecosystems and exert a pervasive influence on “safe operating spaces” for the world’s societal development (Cardinale et al., 2012; Steffen et al., 2015).

Invasive species further exacerbate environmental pressures, constrain the production of food and other agricultural commodities, and disrupt ecosystem functioning – thus undermining efforts to reach the SDGs (Bradshaw et al., 2016; Paini et al., 2017). Regularly tied to the liberalized trade of agricultural produce, invasive species inflict substantial economic losses globally (Pimentel et al., 2001; Bradshaw et al., 2016) and place a disproportionate burden on developing economies and biodiversity-rich tropical settings (Early et al., 2016). Though invasive species have been well-studied through an ecological lens and their impacts are sporadically expressed in monetary terms, their broader (long-term) effects on human well-being and livelihoods have received less attention (Jones, 2017; Shackleton et al., 2019).

Ecosystem alteration, loss of ecological resilience and the appearance of invasive species impact human health in myriad ways (Myers et al., 2013; Sandifer et al., 2015). Plant pathogen invasion in genetically-uniform crops can trigger famine, mass migration and socioeconomic upheaval, as illustrated by the role of fungal blight in the 1845 Irish Potato famine (Cox, 1978). Non-native human pathogens or disease-carrying mosquitoes pose real public health risks (Ricciardi et al., 2011; Medlock et al., 2012), which can be inflated by land-use change or environmental pollution (Myers et al., 2013). In agri-food systems, biodiversity decline compromises food provisioning, downgrades nutritional value of harvested produce and affects welfare (Potts et al., 2010), and persistent anthropogenic pressure on ecosystem services can precipitate a collapse of entire civilizations (Mottesharei et al., 2014). These kinds of non-monetary assessments accentuate the contribution of nature to societal well-being (Daily et al., 2009), and wait to be broadened to other systems, livelihood and vulnerability contexts (Pretty et al., 2018).

One notorious invasive species is the cassava mealybug, *Phenacoccus manihoti* (Hemiptera: Pseudococcidae); a sap-feeding insect that arrived in Africa during the mid-1970s. Causing yield reductions of 80% and regularly leading to total crop loss, *P. manihoti* rapidly spread across Africa’s cassava belt and impacted crop production in 26 countries (Herren & Neuenschwander, 1991; Fig. 1). Cassava, *Manihot esculenta* Crantz (Euphorbiaceae), is a major food staple and a vital source of carbohydrates for local under-privileged populations (Jarvis et al., 2012). Though *P. manihoti* certainly compromised food security at a continental scale, only anecdotal accounts exist of mealybug-induced famine e.g., in northwestern Zambia (Hansen et al., 1994). The invasive mealybug was permanently suppressed through the 1981 introduction of a host-specific parasitic wasp *Anagyrus lopezi* (Hymenoptera: Encyrtidae) from the Neotropics. This biological control (BC) effort permitted a total yield recovery and generated long-term economic benefits that currently surpass US $120 billion (Herren and Neuenschwander, 1991; Zeddies et al., 2001; Raitzer & Kelley, 2008). Aside from these first-order assessments of economic impacts, there has been no comprehensive evaluation of how agro-ecological imbalance (i.e., *P. manihoti* invasion) and its ecologically-centered restoration affected human capital dimensions of African livelihoods.

**Figure 1.**
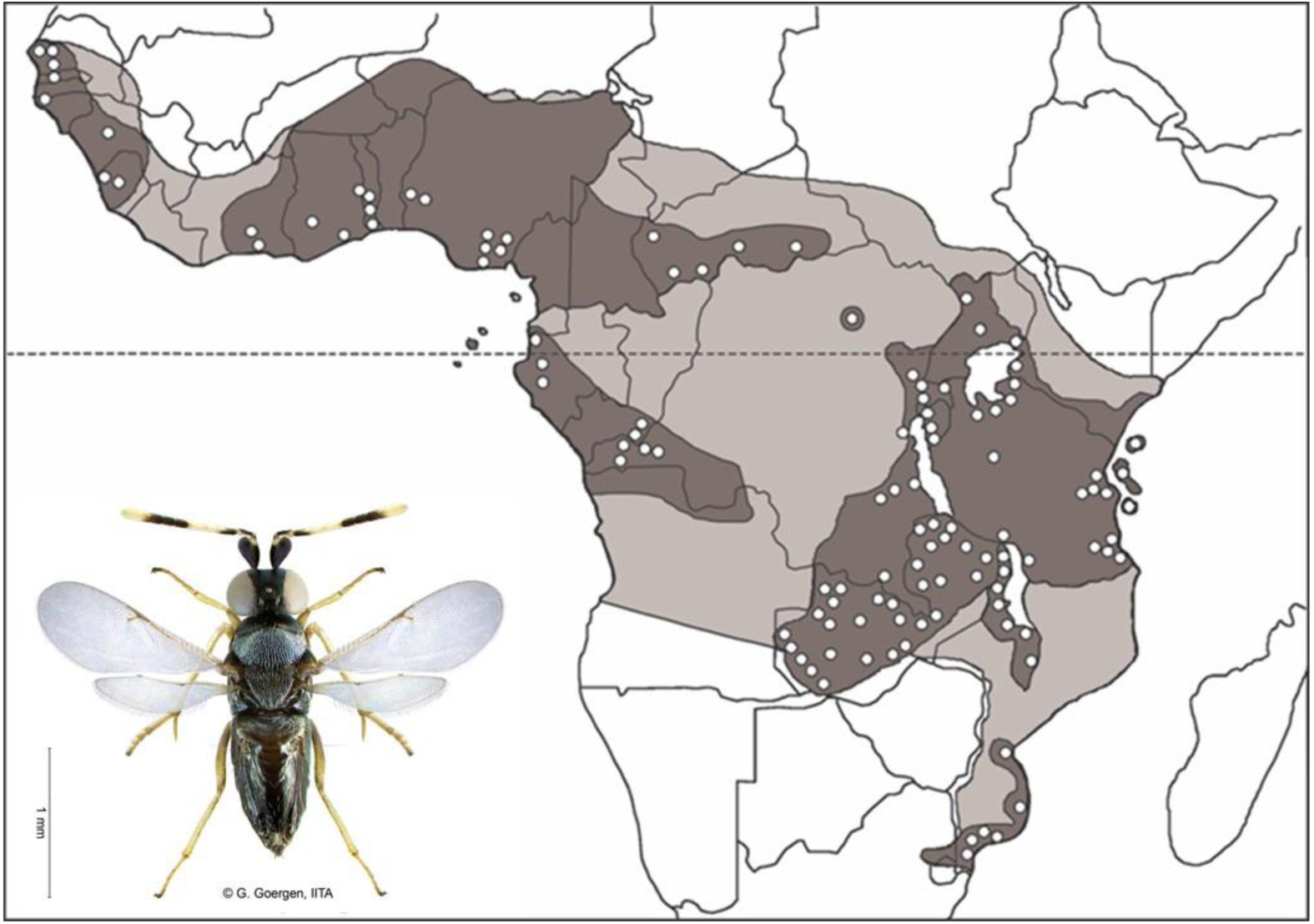
Approximate invasion range of the cassava mealybug (*Phenacoccus manihoti*; light grey area) and the associated distribution of its specialist parasitoid *Anagyrus lopezi* (dark grey area, reflecting spatial spread during the 1990s) across sub-Saharan Africa. Locations of parasitoid releases -as carried out between 1981 and 1995-are indicated as white dots on the map, with one single dot regularly representing multiple locales. Field work after 1995 has confirmed that *A. lopezi* controls cassava mealybug also in the light grey areas. Photo credits of *A. lopezi*: G. Goergen (International Institute for Tropical Agriculture, IITA).

Here, we draw upon historical invasion records, crop production statistics and human population demographics to quantify the extent to which invasive species attack and its ensuing BC impact food availability and human wellbeing in 18 different countries of sub-Saharan Africa. Using demographic metrics as proxies of wellbeing, we empirically assess how 1) mealybug-induced food system collapse triggers a reduction in birth rate and deceleration in population growth; 2) the *A. lopezi* release alleviates nutritional deprivation and its effects on livelihood security and human health; and 3) upsets in biodiversity-mediated ecosystem services (i.e., natural BC) have protracted effects on key livelihood assets. Our work uncovers the extent to which agro-ecological imbalance jeopardizes human well-being over extensive geographical areas and prolonged time periods, and how a restoration of agro-ecosystem health benefits overall human capital.

## Results

### Cassava production shocks and yield recovery

Across sub-Saharan Africa, the mealybug invasion coincided with a max. 18.1 ± 29.4% decline in cassava root yield and a 17.6 ± 28.3% drop in aggregate production over 1-11 years (Table 1). Following parasitoid introduction, a 28.1 ± 34.5% recovery of yield and a 48.3 ± 50.7% increase in production were recorded over variable time periods. Important inter-country differences are observed between the successive time periods (i.e., pre-invasion, post-invasion and post-introduction), for both root yield and aggregate production (Table 1). Maximum pest-induced shocks in yield and production are recorded for Rwanda (−84.3%) and Senegal (−86.7%), respectively. Following *A. lopezi* introduction, the largest recovery of these respective production parameters was reported for Togo (+ 113.5%) and Senegal (+ 208.0%) (Table 1).

**Table 1.** Cassava production shocks and yield recovery following to the *P. manihoti* invasion and ensuing *A. lopezi* introduction. For different cassava production parameters, percent change is computed at a country level either between the five years pre-invasion (averaged) and the post-invasion minima, and between the post-invasion minimum and averages for the max. 10-year parasitoid-induced recovery phase. To account for in-country *A. lopezi* establishment, a 5-year time period is included in the latter. All analyses are conducted over a 1965-1995 window.

| Country | % cassava in savanna | Pest-induced shocks (% decline of pre-invasion baseline) |  | Parasitoid-mediated recovery (% increase of invasion baseline) |  |
| --- | --- | --- | --- | --- | --- |
|  |  | Production | Yield | Production | Yield |
| Congo, Dem. Rep. | 45 | + 5.44 | - 3.40 | + 62.23 | + 18.81 |
| Congo | 60 | + 5.02 | + 3.33 | + 39.70 | + 44.09 |
| CAR | 75 | - 40.91 | - 13.27 | + 5.48 | + 11.46 |
| Gabon | 20 | + 11.84 | - 1.00 | + 17.41 | + 0.08 |
| Benin | 95 | - 8.07 | - 12.27 | + 52.35 | + 23.75 |
| Angola | 18 | - 26.56 | - 6.17 | + 35.67 | + 13.31 |
| Ghana | 67 | - 7.96 | - 16.31 | + 134.95 | + 27.72 |
| Liberia | 10 | - 51.05 | - 15.01 | + 39.43 | + 2.82 |
| Nigeria | 15 | - 7.87 | - 10.61 | + 50.45 | + 16.12 |
| Togo | 95 | - 10.30 | - 79.06 | + 31.48 | + 113.49 |
| Cameroon | 29 | + 5.07 | + 42.83 | + 29.28 | + 12.64 |
| Cote d'Ivoire | 40 | + 13.19 | - 0.66 | + 15.52 | + 1.88 |
| Equatorial Guinea | 0 | + 2.80 | - 14.56 | + 6.94 | + 1.08 |
| Rwanda | 0 | - 53.09 | - 84.33 | + 20.02 | + 106.94 |
| Malawi | 89 | - 47.57 | - 52.53 | + 41.94 | + 24.38 |
| Sierra Leone | 60 | + 8.70 | - 20.16 | + 77.58 | + 69.30 |
| Senegal | 100 | - 86.66 | - 30.30 | + 208.00 | + 19.01 |
| Gambia | -<sup>a</sup> | - 29.08 | - 12.35 | + 0.22 | - 1.67 |
| **Regional average** |  | **- 17.61** | **- 18.10** | **+ 48.26** | **+ 28.07** |

<sup>a</sup>: No data.

### Human demographic impacts of pest attack and biological control

The post-invasion phase was equally typified by declines in birth rate, RNI, fertility rate and increases in adult mortality rate (Table 2). The largest drops in birth rate were recorded for Ghana (9.6%), Togo (10.1%), Rwanda (11.6%) and Senegal (12.30%) (Fig. 2). Upon extrapolating country-level declines, an annual reduction of 377,943 births and a deceleration of RNI by 156,493 was estimated across all the 18 African countries affected by early *P. manihoti* invasions. Over the 10.0 ± 3.6 year-long *P. manihoti* invasion period, this equaled a net loss of 3.26 million births regionally and a total reduction in the natural rate of increase with 838,092 people.

**Table 2.** Shifts in cassava crop production levels and human demographic parameters during the mealybug-invasion and following the *A. lopezi* introduction, for 18 different countries in sub-Saharan Africa. For all parameters, percent change is computed at a country level either between the five years pre-invasion (averaged) and the post-invasion minima, and between the post-invasion minimum and averages for the max. 10-year parasitoid-induced recovery phase. To account for in-country *A. lopezi* establishment, a 5-year time period is included in the latter. All analyses are conducted over a 1965-1995 window.

| Parameter | Pre-invasion baseline | Post-invasion phase (% decline of pre-invasion baseline) | Recovery phase (% increase of invasion baseline) |
| --- | --- | --- | --- |
| Root yield (tonnes/ha) | 6.55 ± 3.59 | -18.10 ± 29.45 | 28.07 ± 34.53 |
| Production ('000 tonnes) | 1,692 ± 3,221 | -17.61 ± 28.30 | 48.26 ± 50.74 |
| Birth rate (/1000 people) | 47.09 ± 4.21 | -5.75 ± 3.93 | -0.17 ± 0.53 |
| Death rate (/1000 people) | 19.11 ± 3.70 | 1.35 ± 22.72 | -12.76 ± 12.45 |
| RNI (/1000 people) | 27.98 ± 3.99 | -4.47 ± 15.41 | 4.66 ± 7.32 |
| Fertility rate (/1000 people) | 6.77 ± 0.75 | -5.47 ± 5.70 | 0.02 ± 1.86 |
| Infant mortality rate (/1000 people) | 126.52 ± 22.55 | -4.84 ± 5.02 | -11.28 ± 10.27 |
| Adult mortality rate (/1000 people) | 404.96 ± 46.72 | 3.73 ± 14.10 | -7.66 ± 9.33 |
| Life expectancy (years) | 46.60 ± 4.62 | 0.08 ± 8.25 | 6.77 ± 7.73 |

**Figure 2.**
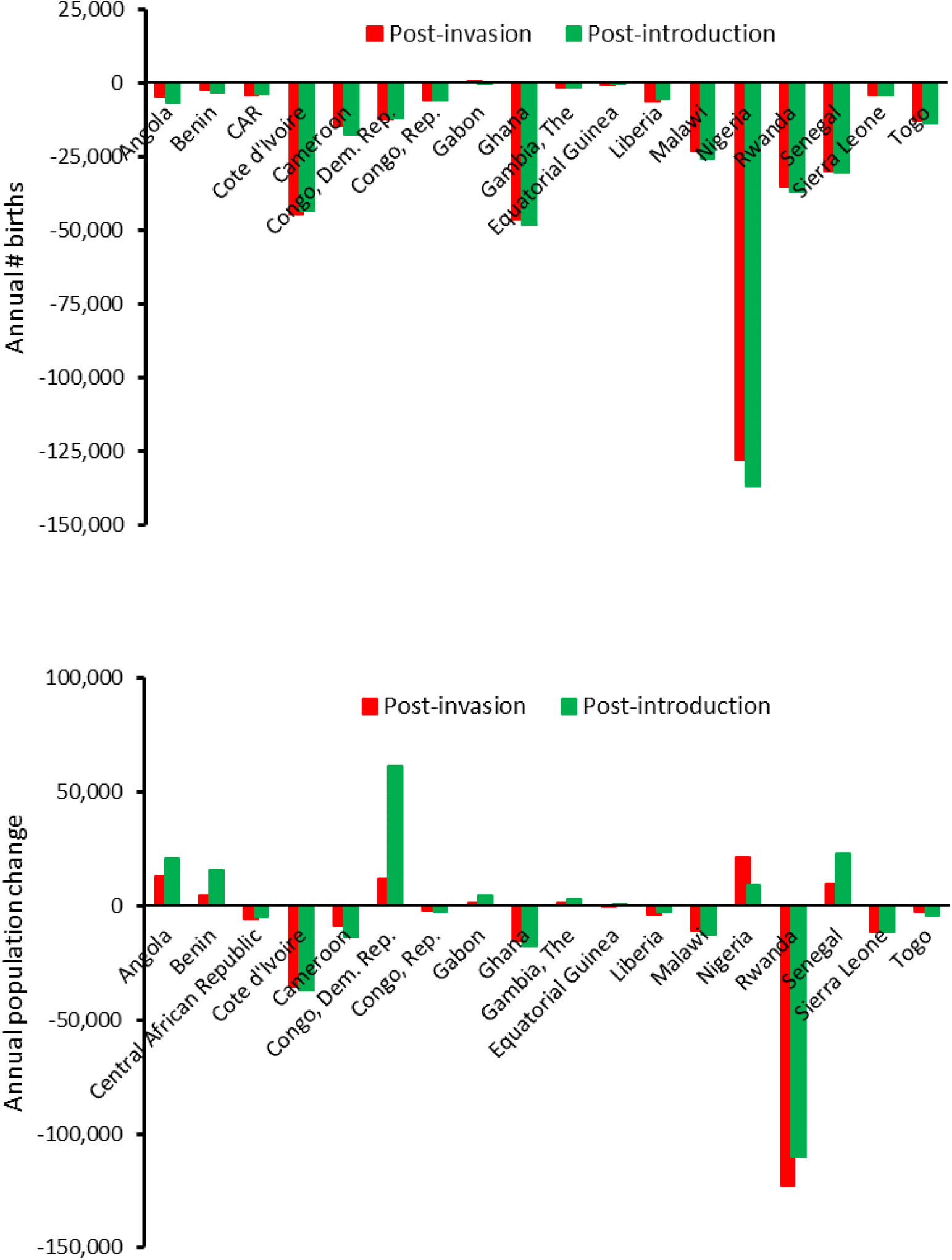
Shifts in the annual number of births and annual population change during the mealybug-invasion and following the *A. lopezi* introduction, for 18 different countries in sub-Saharan Africa. For both demographic parameters, percent annual change is computed at a country-level either between the five years pre-invasion (averaged) and the post-invasion minima, and between the former pre-invasion measure and averages for a max. 10-year parasitoid-induced recovery phase. To account for in-country *A. lopezi* establishment, a 5-year time period is included in the latter. All analyses are conducted over a 1965-1995 window.

On the other hand, restorations in RNI, fertility rate and life expectancy were recorded during the post-introduction phase, while death rate, infant mortality and adult mortality were lowered. Paired-sample tests indicate country-specific changes in birth rate (t= −5.956, df= 17, *p*= 0.000), death rate (t= 2.589, df= 17, *p*= 0.019), RNI (t= −2.273, df= 17, *p*= 0.036), infant mortality (t= 2.262, df= 15, *p*= 0.039), adult mortality (t= 3.769, df= 17, *p*= 0.002), fertility rate (t= −4.278, df= 17, *p*= 0.001), life expectancy (t= −3.096, df= 17, *p*= 0.007) between both post-invasion and post-introduction periods. When computing post-introduction shifts using averaged post-invasion data instead of minima, significant changes were only recorded for birth rate (t= −4.506, df= 17, *p*< 0.001), adult mortality (t= 2.158, df= 17, *p*= 0.046), fertility rate (t= −2.120, df= 17, *p*= 0.049).

During the post-invasion phase, country-level yield losses were related to changes in birth rate (ANOVA, F_1,16_= 5.692, *p*= 0.030, R2= 0.262), RNI (F_1,16_= 6.150, *p*= 0.025, R2= 0.025), death rate (F_1,16_= 6.414, *p*= 0.022, R2= 0.286), adult mortality (F_1,16_= 4.773, *p*= 0.044, R2= 0.230), fertility (F_1,16_= 4.869, *p*= 0.042, R2= 0.233) and life expectancy (F_1,16_= 4.853, *p*= 0.043, R2= 0.233) (Fig. 3; Table 2). Marginally significant patterns were recorded between cassava production declines and changes in birth rate (F_1,16_= 4.394, *p*= 0.052, R^2^= 0.215). During the post-introduction phase, no significant trends were recorded for yield gain with any of the demographic parameters, while marginally-significant regressions were found between aggregate production and infant mortality (F_1,16_= 4.238, p= 0.056). Significant country-specific effects were recorded of the *P. manihoti* invasion and *A. lopezi* introduction on all crop production and human demographic parameters (Supplementary Table 1). When accounting for temporal trends in cassava production and demographic parameters, statistical significance of those patterns was sustained.

**Figure 3.**
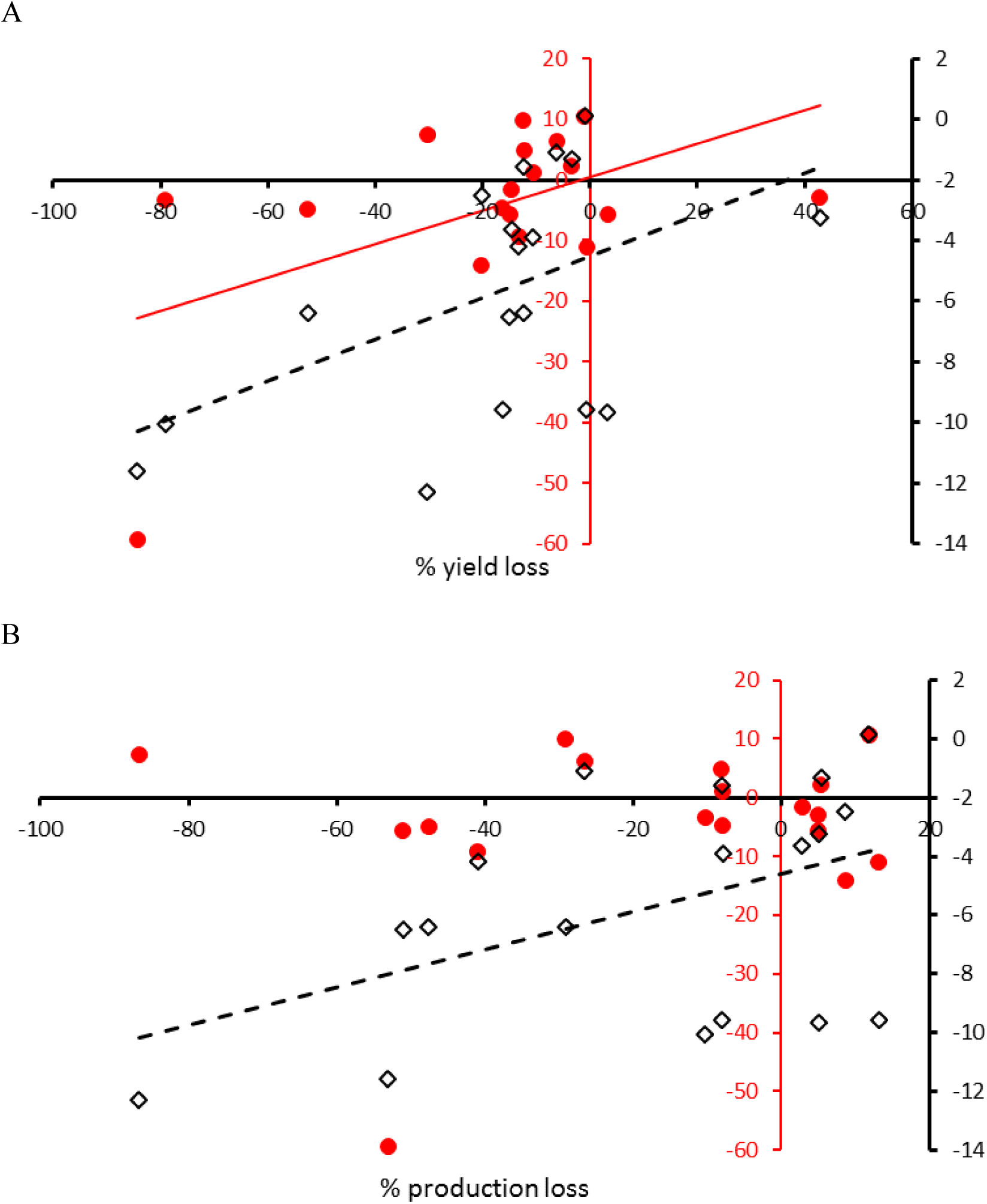
Mealybug-induced shocks in cassava crop yield (A) and production (B) impact different human demographic parameters, as outlined for 18 different countries in sub-Saharan Africa. Graphs depict statistically-significant regression lines for either birth rate (black diamonds, dashed lines) and natural rate of increase (red dots, full line). For each demographic and crop production parameter, percent change is computed at a country level between the five years pre-invasion (averaged) and the post-invasion minima. All analyses are conducted over a 1965-1995 window. Statistical details are covered in the text.

Generalized additive regression modelling allowed year-by-year assessment of the relative impacts of yield shifts and in-country presence of biotic stressors on demographic parameters for two specific events (i.e., post-invasion, post-introduction). During the post-invasion phase, full-factorial models with a time-step and yield shift provided the best goodness-of-fit (Table 3; AIC = 190.84 and 179.99 for birth rate and RNI respectively). Models yielded significant negative impacts of the interaction term, i.e., yield and time-step on both demographic parameters. During the post-introduction phase, full-factorial models with the interaction term of biotic stress and yield-shift provided the best fit (Table 3; AIC = 497.54 and 216.93 for birth rate and RNI respectively). Models revealed a significant impact of the interaction term on both demographic parameters.

**Table 3.** Generalized additive regression models for proportional shifts in rate of natural increase (RNI) and birth rate following the *P. manihoti* invasion (post-invasion) and *A. lopezi* introduction (post-introduction). Explanatory variables include yield shift between pre-invasion values yearly post-invasion values, and in the case of post-introduction analysis, yield shifts between pre-and post-introduction of *A. lopezi*. For post-invasion analysis, an additional variable named “time step” was included, while for post introduction, an additional variable ‘biotic stress’ was included reflecting in-country mealybug or parasitoid presence. Three different regression models were contrasted, for which estimates, corresponding p-values and Cragg Uhler R-squared values are represented.

| Response variable | Model | Predictor variable | Estimate | p-value | Cragg Uhler R squared |
| --- | --- | --- | --- | --- | --- |
| **Post-invasion** |  |  |  |  |  |
| Birth rate | 1 | Time step | - 0.032 | 0.432 | 0.009 |
|  | 2 | Yield shift | 0.328 | 0.00458 ** | 0.076 |
|  | 3 |  |  |  | 0.13 |
|  |  | Time step | - 0.0009 | 0.984 |  |
|  |  | Yield shift | 0.831 | 0.00142 ** |  |
|  |  | Interaction | - 0.218 | 0.04777 * |  |
| RNI | 1 | Time step | 0.09834 | 0.00695 ** | - 0.032 |
|  | 2 | Yield shift | - 0.009052 | 0.942 | - 0.0795 |
|  | 3 |  |  |  | 0.15 |
|  |  | Time step | 0.17744 | 1.6e-05 *** |  |
|  |  | Yield shift | 0.69734 | 0.23259 |  |
|  |  | Interaction | - 0.19520 | 0.000716 *** |  |
| **Post-introduction** |  |  |  |  |  |
| Birth rate | 1 | Biotic stress | 0.5035 | 1.10e-08 *** | 0.15 |
|  | 2 | Yield shift | 0.18 | 0.000282 *** | 0.001 |
|  | 3 |  |  |  | 0.29 |
|  |  | Biotic stress | 0.656 | 6.97e-11 *** |  |
|  |  | Yield shift | 0.454 | 8.01e-13 *** |  |
|  |  | Interaction | - 0.323 | 0.000726 *** |  |
| RNI | 1 | Biotic stress | - 0.009 | 0.936 | - 0.29 |
|  | 2 | Yield shift | 0.22 | 2.00e-16 **** | 0.56 |
|  | 3 |  |  |  | 0.60 |
|  |  | Biotic stress | 0.011 | 0.256 |  |
|  |  | Yield shift | 0.396 | 0.00467 ** |  |
|  |  | Interaction | - 0.208 | 0.0918. |  |

## Discussion

Invasive terrestrial invertebrates cause major economic impacts, with insects inflicting US $70 billion globally in direct costs (Bradshaw et al 2016). Such monetary estimates are conservative, do not fully capture the long-term adverse effects of invaders on biodiversity-mediated ecosystem services, e.g., the US $400 billion annual service of natural biological control (BC; Costanza et al., 1997), and only reveal a fraction of their broad societal impacts. Here, we demonstrate empirically how a crop-damaging insect adds to a multi-country decline of birth rate (−5.8%) and fertility rate (−5.5%), compounded by elevated adult mortality (3.7%) and a leveling life expectancy (0.1%). Through inducing a primary productivity loss of cassava; one of Africa’s main food staples and a valued source of dietary carbohydrates (Jarvis et al., 2012), *P. manihoti* triggered sequential periods of food insecurity and nutritional deprivation along its 1970-1980s invasion path. Mirrored in an annual decline of 380,000 births and an RNI deceleration of 156,500 people, the mealybug caused hardship, deepened poverty and disrupted reproduction plans for an approximate 162 million African citizens over a 10-year period. The 1981 *A. lopezi* release permanently resolved continent-wide mealybug issues (Neuenschwander, 2001), contributed to a 48.3% recovery of total crop output and enabled parallel improvements in multiple demographic indices (Table 1, 2). This compelling case accentuates the social and human health repercussions of ecological upsets in subsistence farming systems, providing lessons for global efforts to resolve food insecurity, mitigate invasive species and safeguard functionality and resilience of (agro-)ecosystems.

Our estimates of parasitoid-mediated yield recovery are in line with earlier estimates of 15-37% under forest and savanna conditions, respectively (Zeddies et al., 2001), and substantially lower than the 50.0-96.6% loss reduction in Ghana’s savanna (Neuenschwander et al., 1989). Yet, the parallel invasion of the spider mite, *Mononvchellus tanajoa* (Bondar) is likely to have inflated yield losses in certain areas (Yaninek et al., 1993). On the other hand, our exclusion of certain countries where *P. manihoti* and *A. lopezi* either arrived simultaneously, multiple invasion instances occurred, ground-truthing wasn’t carried over 1975-1995, or environmental heterogeneity didn’t allow an unambiguous assessment of biotic impacts certainly led to an underestimation of continental-scale BC benefits. Yet, our work confirms previous assessments of the agronomic impacts of *A. lopezi* for e.g., Nigeria or Ghana, and can be a reliable indicator of its ensuing benefits for food security and human well-being especially in western sections of sub-Saharan Africa.

Diagnosing food insecurity is notoriously difficult, and food availability measures *in se* have low predictive accuracy (Sen, 1981; Barrett, 2010). Pest-induced yield loss in a core food staple directly degrades provisioning services and causes food system failure, with distinct livelihood impacts. Extended periods of food shortage and mal-nutrition regularly progress into famine and population loss (Scrimshaw, 1987). Though literature records confirm mealybug-induced famine in certain geographies (Hansen, 1994), our reports of lowered birth rate, increased mortality and slowing population growth signal a long-term, severe nutritional deprivation (Kane, 1987; Scrimshaw, 1987). Famine syndromes are also reflected in a lowered birthweight, increased stillbirths or premature births, and higher rates of voluntary birth control - though these events likely went unrecorded in disrupted rural populations during the study period. In subsistence systems, crop failure causes loss of direct food entitlement among peasants but may also drive up food prices and generate chronic ‘poverty traps’ (Sen, 1981; Kane, 1987; Swinnen & Squicciarini, 2012). Nutritional deprivation further brings about social upheaval (e.g., disrupted family structure, delayed marriages, migration), leads to -often lagged-psychological change or lowers resistance of malnourished populations to disease attack (Cox, 1978; Painter et al., 2005). Hence, aside from the 3.2 million foregone births over the span of the mealybug invasion, other human and social dimensions of African livelihoods certainly were affected.

Given the distinctive invasive species impacts on livelihoods and human well-being, proactive mitigation strategies are essential and agricultural ecosystems will sufficiently resilient to sustain delivery of multiple ecosystem services under a range of external stressors (Ricciardi et al., 2011; Jones, 2017; Shackleton et al., 2019; Early et al., 2016; Pretty et al., 2018). The framework of food systems lends itself to fuse ecological facets of global change -e.g., species invasion-with social or human aspects, as to optimally interpret their societal outcomes (Ericksen, 2008; Ingram, 2011). As basis for many of today’s food systems, conventional agriculture seldom provides simultaneous positive outcomes for ecosystem service provisioning and human well-being (Garibaldi et al., 2017; Rasmussen et al., 2018), and often allows biotic shocks to proliferate and cascade e.g., into socio-economic domains (Wyckhuys et al., 2018). A stabilization or strengthening of the ecological foundation of food systems can bolster resilience and dampen such ill-favored externalities (Cox, 1978; Fraser et al., 2005). One way to achieve this is through ecological intensification: an integrated set of interventions that harness ecosystem services - including ‘biotic resistance’ to pest invasion, conserve crop yields and often enhance farm profit (Bommarco et al., 2013; Garbach et al., 2017). Organic agriculture consistently spawns ‘win-win’ outcomes for natural and human capital, and a targeted redesign of conventional farming systems can also help achieve those goals (Reganold & Wachter, 2016; Pretty et al., 2018; Eyhorn et al., 2019). Such transformation can be enabled by consciously prioritizing resource-conserving biodiversity-based approaches, and by integrating agro-ecological metrics in government decision-making, food crisis vulnerability diagnostics or along the food value chain (Gordon et al., 2017; Sukdev, 2018).

As an environmentally-sound approach to pest management, BC requires a major reassessment by all those responsible for achieving a sustainable planet (Bale et al., 2008). Since the late 1800s, BC has enabled the long-term suppression of over 200 invasive insect species at benefit:cost ratios often >1,000:1 (Naranjo et al., 2015; Heimpel & Mills, 2017). Modern biological control, centered on a careful selection of a specialized natural enemy (e.g., the monophagous parasitoid *A. lopezi*) effectively resolves invasive species issues. Also, as a globally-important ecosystem service, natural BC underpins resilience of agro-ecosystems (Costanza et al. 1997). Ensuring durable, cost-effective control of crop pests, BC delivers sustainable human health, economic and environmental outcomes (Bale et al., 2008; Naranjo et al., 2015); a valuable service for the resource-poor smallholders that secure global food security. While a scientifically-guided introduction of natural enemies can restore ecological balance and diverse, extra-field habitats usually support BC, a field-level reliance on biodiversity-friendly practices is pivotal to the effective conservation of agro-ecosystems’ natural functionality (Landis et al., 2000; Karp et al., 2018). Disruptive interventions, e.g., misuse of synthetic pesticides adversely impact beneficial insects, accelerate pest proliferation and nullify BC benefits (Geiger et al., 2010; Lundgren and Fausti, 2015), at a risk of inflating crop damage substantially (Losey & Vaughan, 2006). Instead, our study illuminates how BC boosts on-farm functional biodiversity, lifts the carrying capacity of subsistence farming systems across Africa (Mottesharrei et al., 2014), and generates concrete societal dividends.

Invasive species attack, rapid biodiversity depletion and progressive ecosystem simplification undermine multiple of the UN Sustainable Development Goals (Dobson et al., 2006; Cardinale et al., 2012; Bradshaw et al., 2016; Oliver et al., 2015). In order to meet the SDG implementation targets, integrative approaches between social and natural sciences and a deliberate recognition of inter-sectoral linkages are needed (Brondizio et al., 2016; Stafford-Smith et al., 2017). Building a solid human capital foundation for ecologically-intensified farming, we enclose pest-induced nutritional deprivation (and cascading livelihood impacts) within an overarching food systems framework. Our work accentuates how biodiversity-based techniques can help meet food security needs by fortifying agro-ecosystem functionality (Godfray et al., 2010; Stephens et al., 2018), and thus generate ample spillover benefits for collective societal welfare (Rasmussen et al., 2018; Pretty et al., 2018). If the current biodiversity crisis is a warning sign of impending agro-ecological imbalance, swift and deliberate action can prevent food-related socio-economic hardship from becoming a recurrent feature of our uncertain future.

## Data Availability

All data underlying the analyses in this manuscript will be made available through Dryad Digital Repository.

## Acknowledgements

The development of this manuscript and its underlying research received no noteworthy funding.

## Author contributions

KAGW conceived and designed the experiments; KAGW performed trials and collected the data; KAGW and DDB analyzed the data; KAGW, DDB, JP and PN co-wrote the paper.

## Materials and Methods

### Data

Invasion history for *P. manihoti* and associated country-level introductions of *A. lopezi* -as conducted during 1981-1995-were obtained from Herren et al. (1987), Neuenschwander (2001) and Zeddies et al. (2001). Country-specific patterns of cassava production (harvested area, ha; tonnes) and fresh root yield (tonnes/ha) were obtained for all mealybug-invaded countries through the FAO STAT database (http://www.fao.org/faostat/). Historical country-level records for a range of human demographic parameters were accessed through the World Bank Open Data portal (https://data.worldbank.org/). More specifically, country-specific data-sets were obtained for birth rate, death rate, fertility rate, rate of natural increase (RNI), infant mortality and adult mortality. Analyses centered upon a total of 18 different African countries that were affected in the early stages of the mealybug invasion, primarily including countries in West and Central Africa (Herren et al., 1987). These constitute a sub-set of the 27 African nations that were impacted by *P. manihoti* (Zeddies et al., 2001).

### Cassava production shocks and yield recovery

To assess changes in cassava production following the *P. manihoti* invasion and the *A. lopezi* introduction, we examined temporal shifts in country-level root yield and aggregate production. More specifically, we contrasted yield and production trends over three different time periods: a 5-year pre-invasion period, a post-invasion period of variable duration, and a post-introduction period that followed the first in-country detection of the parasitoid (see Zeddies et al., 2001). Over a three-year time period, *A. lopezi* successfully established in 70% fields of a given country (Herren et al., 1987), and we assume that its biological control impacts become well-apparent after 5 years following its in-country release.

More specifically, we detailed changes in aggregate cassava production (tonnes) and yield between the above time periods - with assessments spanning a 1965-1995 window. We used a general linear model (GLM) to detect statistical differences in yield and aggregate output between successive time periods, using as explanatory variables country, a dummy variable reflecting in-country mealybug and parasitoid presence, and the respective interaction term. Accounting for broader temporal trends in cassava production and the possibility of model over-fit, we equally carried out a GLM using the residuals of a general linear regression analysis of yield and production vs. time. To meet assumptions of normality and heteroscedasticity, data were subject to rank-based inversed normal transformation. All statistical analyses were conducted using SPSS (PASW Statistics 18).

### Country-specific impacts of pest attack and biological control on human demography

Over the above time periods, we characterized temporal trends in multiple human demographic parameters. For each parameter, we also compared across all countries proportional shifts over the post-invasion and the post-introduction phase using a paired-samples t test. Proportional changes were computed at a country level either between the five years pre-invasion (averaged) and the post-invasion minima, and between the post-invasion minimum and averages for the max. 10-year parasitoid-induced recovery phase. As such, we captured a worst-case scenario for either the post-invasion or post-introduction phase. Selected analyses were re-run using proportional changes that were based upon averaged post-invasion values (instead of minima).

Next, we related either *P. manihoti*-induced declines or the *A. lopezi*-assisted recovery in cassava production to country-level changes in human demographics. More specifically, we conducted analyses to assess production-related impacts on the following demographic parameters: birth rate, fertility rate, RNI, infant mortality and adult mortality. First, linear regression was carried out to relate proportional changes in cassava root yield and production (for either the post-invasion period, and the post-introduction phase) to proportional changes in birth rate and RNI over these respective time periods. Linear regression analysis covered all 18 mealybug-invaded countries, examining patterns over a 1965-1995 window.

Second, we employed a general linear model (GLM) to detect statistical differences in human demographic parameters, using as explanatory variables country, a dummy variable reflecting in-country mealybug and parasitoid presence, and the respective interaction term. Considering an eventual generalized trend in human demographics across sub-Saharan Africa and to control for model over-fit, we equally carried out GLM using the residuals of a linear regression analysis of different demographic parameters with time (i.e., year).

### Multi-country, year-by-year impacts of pest attack and biological control

In addition to using pre-and post-invasion average values and post-invasion minima for regression analysis, we also conducted a generalized additive regression model, using year-by-year values of root yields, birth rate and RNI, in order to elucidate impacts of yield shifts across countries. For regression analysis, proportional shifts in root yield, RNI and birth rate were calculated by differencing yearly post-invasion and post-introduction values of these parameters with 5-year averaged values pre-invasion. For post-invasion regression analysis, root yield, RNI and birth rate shifts corresponded to differenced values, between 5-year averaged values pre-invasion, with year-by-year value post invasion. A dummy variable, time-step, was constructed reflecting the number of years since invasion. In the case of post-introduction regression analysis, yield, birth rate and RNI shifts, also included differenced values between 5-year averaged values pre-invasion to those year-by-year values post-introduction, in addition to differenced values of 5-year averaged values pre-invasion, with year-by-year value post-invasion. Additionally, in the regression analysis of post-introduction phase, a dummy variable, biotic stress representing in-country mealybug and/or parasitoid presence (i.e., post-invasion and post-*A. lopezi* introduction) was used. Sets of regression equations were computed for either RNI or birth rate as outcome variables, and time step, proportional yield shift for post-invasion analysis, and biotic stress and proportional yield shift for post-introduction analysis, as explanatory variables. Due to the complex distribution of the shift values, a Generalized Additive Model for Location, Scale and Shape (GAMLSS) approach was used for regression analysis. For each set, three regression models were compared, i.e. either using the dummy variable or yield shift as explanatory variable and a full factorial (both previous variables plus interaction term). Goodness-of-fit of models was assessed by comparing the three sets of regression equations for both post-invasion and post-introduction phase, using the global Akaike information criterion score (AIC), CraggUhler R-squared metric and correlation metric between predicted and actual values of the demographic parameters (i.e., birth rate and RNI) as response variables. The fitdist approach was used to identify and fit the distribution for the response variables. Regression analysis was performed using the GAMLSS package in R (v 3.4.1) (Stasinopoulos et al., 2017).

